# Regulation of mitochondrial calcium uniporter expression and calcium signalling by lncRNA *Tug1* in cardiomyocytes

**DOI:** 10.1101/2023.07.22.550175

**Authors:** Adam J. Trewin, Kate L. Weeks, Glenn. D. Wadley, Séverine Lamon

**Affiliations:** Institute for Physical Activity and Nutrition, and School of Exercise and Nutrition Sciences, Deakin University, Geelong, Australia; Department of Anatomy and Physiology, University of Melbourne, Australia; Baker Department of Cardiometabolic Health, University of Melbourne, Australia

**Keywords:** Cardiac, mitochondria, transcriptome, non-coding RNA, calcium signalling

## Abstract

Cardiomyocyte calcium homeostasis is a tightly regulated process. The mitochondrial calcium uniporter (MCU) complex can buffer elevated cytosolic Ca^2+^ levels and consists of pore-forming proteins including MCU, and various regulatory proteins such as mitochondrial calcium uptake proteins 1 and 2 (MICU1/2). The stoichiometry of these proteins influences the sensitivity to Ca^2+^ and activity of the complex. However, the factors that regulate their gene expression remain incompletely understood. Long non-coding RNAs (lncRNAs) regulate gene expression through various mechanisms, and we recently found that the lncRNA *Tug1* increased the expression of *Mcu* and associated genes. To further explore this, we performed antisense LNA knockdown of *Tug1* (*Tug1* KD) in H9c2 rat cardiomyocytes. *Tug1* KD increased MCU protein expression, yet pyruvate dehydrogenase dephosphorylation, which is indicative of mitochondrial Ca^2+^ uptake was not enhanced. However, RNA-seq revealed that *Tug1* KD increased *Mcu* along with differential expression of >1000 genes including many related to Ca^2+^ regulation pathways in the heart. To understand the effect of this on Ca^2+^ signalling, we measured phosphorylation of Ca^2+^/calmodulin-dependent protein kinase II (CaMKII) and its downstream target cAMP Response Element-Binding protein (CREB), a transcription factor known to drive *Mcu* gene expression. In response a Ca^2+^ stimulus, the increase in CaMKII and CREB phosphorylation was attenuated by *Tug1* KD. Inhibition of CaMKII, but not CREB, partially prevented the *Tug1* KD- mediated increase in *Mcu*. Together, these data suggest that *Tug1* modulates MCU expression via a mechanism involving CaMKII and regulates cardiomyocyte Ca^2+^ signalling which could have important implications for cardiac function.

## INTRODUCTION

Calcium is an essential secondary messenger for signalling, excitation contraction and energy homeostasis in the cardiomyocyte (1). In systole, Ca^2+^ entry via voltage-gated L-type Ca^2+^ channels (LTCC, also known as the dihydropyridine receptor, encoded by *Cacna1c*), triggers the ryanodine receptor 2 (*Ryr2*) to release Ca^2+^ from the sarcoplasmic reticulum (SR). The resulting increase in cytosolic Ca^2+^ leads to excitation-contraction coupling and contraction of the sarcomere. SR Ca^2+^ stores are then replenished during diastole by Ca^2+^-ATPase 2a (SERCa2, *Atp2a2*) and efflux from the cell occurs through the Na^+^/Ca^2+^-exchanger (NCX) and the plasmalemmal Ca^2+^-ATPase (PMCA). Ca^2+^ fluxes are sensed by several signalling pathways including Ca^2+^/calmodulin dependent kinase II (CaMKII), which acutely regulates various ion channels and Ca^2+^ handling proteins through post translational modifications, as well as regulating gene expression via several transcription factors including CREB (2). Throughout these rapid fluxes as well as longer term shifts in steady state Ca^2+^ levels, elevated cytosolic Ca^2+^ levels are buffered by mitochondria via the mitochondrial calcium uniporter (MCU) complex.

Mitochondria play important roles in a range of biological processes beyond ATP synthesis, including maintenance of cellular Ca^2+^ homeostasis via the MCU. Mitochondrial Ca^2+^ uptake via the MCU sequesters excess cytosolic Ca^2+^, but overload can lead to opening of the mitochondrial permeability transition pore, induction of apoptosis and cell death (3). On the other hand, insufficient mitochondrial Ca^2+^ uptake impairs TCA activity, disrupting cellular bioenergetics and ultimately leading to myocyte dysfunction (3). Therefore, optimal mitochondrial function relies on the fine regulation of calcium homeostasis (4). The MCU complex consists of pore-forming proteins MCU, MCUb, EMRE as well as regulatory proteins mitochondrial calcium uptake proteins 1 to 3 (MICU1-3) (5). The stoichiometry of these proteins influences the sensitivity to Ca^2+^ and activity of the complex (6). However, the upstream factors that regulate their expression remain incompletely understood.

Long non-coding RNAs (lncRNAs) can affect gene expression through various mechanisms that regulate transcription processes (7). We recently identified the lncRNA *Tug1* as a modulator of mitochondrial and myogenic transcriptional pathways in skeletal muscle (8). In particular, we found that *Tug1* affected the expression of genes that encode MCU complex proteins. This may have implications for cardiomyocyte function, since dysregulation of Ca^2+^ is central to the pathophysiology in cardiac arrythmias, cardiomyopathies and heart failure (9-11). A greater understanding of the factors that regulate MCU expression is needed to open new avenues of investigation into therapies aimed at improving or preventing heart disease. Therefore, in this study, we aimed to determine whether *Tug1* plays a role in cardiomyocyte Ca^2+^ signalling and what are the potential mechanism(s) involved. We report that *Tug1* regulates MCU expression in cardiomyocytes, which occurs in part via CaMKII, and that this has pleiotropic effects on the cardiomyocyte transcriptome.

## MATERIALS AND METHODS

### Cell culture

H9c2 rat cardiomyoblasts (RRID: CVCL_ 0286) were sourced from CellBank Australia (Westmead, NSW, Australia). Cells were cultured at 37°C with 5% CO_2_ in a humidified incubator in proliferation media consisting of Dulbecco’s Modified Eagle Medium (DMEM; Gibco #11995-065) supplemented with 1% v/v penicillin-streptomycin (Gibco #15140-122) and 10% v/v FBS (Gibco #A3161001). Cells were grown in T75 flasks to ∼80% confluence then washed with sterile PBS and detached (Gibco TrypLE Express #LTS12604021) for passaging into 6- or 12-well plates. After 24 h in proliferation media, cells were differentiated for 5 days in media consisting of DMEM (Gibco #11995-065) supplemented with 1% v/v penicillin-streptomycin (Gibco #15140-122), 1% v/v FBS (Gibco #A3161001) and 1 µM retinoic acid (Sigma R2625) which was added to media daily and protected from light. For some experiments, stock solutions of ionomycin (Thermo I24222), CaMKII inhibitor XII (Calbiochem #208923) and CREB inhibitor 666-15 (Millipore #5.38341) were each prepared in sterile DMSO then diluted in DMEM to add to wells to the indicated final concentration.

### Gene silencing *in vitro*

The antisense locked nucleic acid (LNA) GapmeR (#339515, Qiagen) sequence for the *Tug1* LNA was (5’-3’) ATTCAGTAGACAGCTA (Cat # LG00798122-DDA), and control LNA was (5’-3’) GCTCCCTTCAATCCAA (cat # LG00000001-DDC). Lyophilised LNAs were first resuspended in nuclease free water to make a stock solution (50 µM). Lipofectamine® 2000 transfection reagent (Thermo Fisher Scientific) was diluted in serum-free DMEM (7% v/v) and in a separate tube, LNAs were diluted in serum-free DMEM. The diluted LNAs and diluted Lipofectamine were then mixed 1:1 and incubated at room temperature for 15 min to form a final transfection mix. Transfection mix was then added to each well to a final LNA concentration of 25 *n*M. LNA transfection was performed for 24 h prior to subsequent experimental measurement unless otherwise stated.

### RNA isolation

Media was aspirated then cells were immediately lysed in 900 µL TRI-reagent (#15599018 Ambion) with homogenisation achieved by pipette mixing. Lysates were then frozen at −80°C until RNA isolation. TRI-reagent lysates were thawed on ice and centrifuged at 10,000 x g for 10 min at 4°C. An aliquot of the supernatant was transferred to a new tube and mixed with an equal volume of 96% ethanol, then transferred to a spin column for RNA purification with on-column DNase-I treatment, as per manufacturer’s directions (#R2052 Direct-Zol RNA Miniprep, Zymo Research). RNA concentration and purity (260:280 nm: >2.0) was determined with a spectrophotometer (NanoDrop 1000, Thermo Fisher Scientific).

### Reverse transcription and quantitative PCR (RT-qPCR)

Total RNA was reverse transcribed to first-strand cDNA in a 20 µL reaction along with no-template and no-Rtase controls (#4368814 Applied Biosystems). Quantitative PCR was performed in triplicate (Agilent AriaMX G8830A) on 4 ng of cDNA samples (along with a no-template and a no-RT control) in a 10 µL reaction consisting of SYBR green master mix (#4367659 Applied Biosystems) with forward and reverse primers as follows: *Gapdh* 5’-AAGGTCGGTGTGAACGGATTT-3’ and 5’- CCACTTTGTCACAAGAGAAGGC-3’; *Tug1* 5’-AAGTGAACTACGTCCCGTGC-3’ and 5’- CCAGGTCTGTAGGCTGATGG-3’; *Mcu* 5’-TTGTGCCCTCTGATGACGTG-3’ and 5’- ACGGAGTCCGAGATAGGCTT-3’; *Nr4a3* 5’-CAGCAGCTGTGACTCTCCC-3’ and 5’- CGCAGTGGGCTTTGGGT-3’. Thermal conditions for qPCR were an initial 10 min activation step at 95°C and then 40 cycles of 15 s at 95°C denaturing and 60 s anneal/extend at 55-60°C. A dissociation curve was performed to confirm the amplification of a single product. Quantification cycle (C_q_) thresholds were calculated using software (Agilent Aria v1.5) and normalised to a reference gene (*Gapdh*) using 2^-(ΔΔCq).

### RNA sequencing and bioinformatics

RNA concentration was confirmed fluorometrically (Qubit, Thermo Fisher) and RNA integrity number (≥9) was determined by electrophoresis (Tapestation, Agilent). Stranded libraries were prepared from 500 ng total RNA input with poly-A+ selection (TruSeq Stranded mRNA, Illumina). Final library size and concentration were assessed (Qubit, Thermo Fisher and TapeStation, Agilent). Libraries were sequenced with 150bp paired end reads on the Illumina NovaSeq 6000 platform (Macrogen Oceania), generating at least 40 million reads per sample.

Reads underwent quality check with FastQC v0.11.7. Raw read quality filtering and adapter trimming was conducted with fastp v0.14.1 (12) (auto detect of adapters, trim_tail=1, poly_g_min_len=1, min phred quality 20, min length 36, u 90). Filtered reads were aligned to the mouse reference genome (*Mus musculus*, Ensembl version GRCm39.104) using STAR v2.5 (13) in 2-pass mode. Gene expression was quantified at the gene level using HTseq-counts and collated into a m x n matrix. Analysis of differential expression was performed on genes with ≥ 1 count per million (CPM) in at least 3 out of the 6 libraries using Voom/Limma in Degust v 4.1.5 (14). Transcripts with a false discovery rate (FDR) <0.05 were considered differentially expressed. Gene pathway analyses were performed in iDEP v0.96 (15). Transcription factor enrichment was determined from DE gene lists using X2K (16).

### Western blot analysis

Cells were washed in PBS then homogenised on ice with RIPA buffer (#20-188, Millipore) containing protease/phosphatase inhibitors with EDTA (#78442 HALT, Thermo). Samples were centrifuged at 10,000 x g for 5 min at 4°C. Protein concentration was determined by BCA assay (#23225, Thermo Scientific). Samples were mixed with reducing loading buffer (Laemmli sample buffer and 2.5% 2- mercaptoethanol). Samples (∼10 µg protein), molecular weight marker (Bio-Rad #161-0373) and pooled sample (used to generate a standard curve) were loaded on each gel (4-15%, Bio-Rad #5678085). Protein was separated by electrophoresis at 40 V for 30 min then 120 V for 60 min. The gel was imaged for stain-free total protein (Chemi Doc XR+, Bio-Rad) with ImageLab software (Image Lab v6, Bio-Rad) then proteins were transferred (Turbo Transfer system, Bio-Rad) to a methanol-saturated polyvinylidene difluoride (PVDF) membrane (Millipore Immobilon FL 0.45 µm #IPFL00010). The membrane was washed with PBS, then blocked for 1 hour (Li-Cor Intercept PBS blocking buffer). The membrane was then incubated with primary antibody diluted 1:1000 in blocking buffer with 0.2% v/v Tween-20 overnight at 4°C. Primary antibodies used were as follows: MCU (CST 14997), Phospho- Pyruvate Dehydrogenase α1 Ser293 (CST 31866), Phospho-CaMKII Thr286 (CST 12716), Phospho-CREB Ser133 (CST 9196). Blots were then PBS-T washed and incubated in anti-rabbit or anti-mouse IgG Dylight® 800 nm (CST 5151S and 5257S, respectively) secondary antibodies at 1:20,000 in blocking buffer containing 0.2% Tween-20 and 0.01% SDS for 1 hour at room temperature. Images were acquired (Odyssey® Infrared Imaging System, Licor, Lincoln, NE, USA) and blot densitometry performed using software (Odyssey v2.1, Licor). Blot density and stain-free total protein density for each sample was calculated by linear regression using the standard curve constructed from the pooled sample loaded on each gel (17). Calculated blot density values were then normalized to total protein (stain free).

### Statistical analysis

All statistical analyses (except RNA sequencing bioinformatics) were conducted using GraphPad Prism (v8.0; CA, USA). Data are presented as mean(SD) and analysed by two-way ANOVA with adjustment for multiple comparisons as described in the figure legend.

## RESULTS

*Tug1 knockdown increases MCU expression and alters a Ca*^*2+*^*-sensitive mitochondrial pyruvate dehydrogenase response*.

We recently reported that knockdown (KD) of the lncRNA *Tug1* led to increased *Mcu* gene expression (which encodes the main pore-forming protein of the MCU complex) in C2C12 mouse myotubes and in H9c2 cardiomyocytes (8). To further explore the implications of this finding in the cardiac setting, H9c2 cardiomyocytes were transfected with antisense locked nucleic acid (LNA) oligos for 24 hours to knock down (KD) *Tug1* to ∼40% of the control LNA, as determined by qPCR (Figure 1A). There was a concomitant increase in MCU protein abundance over 24 and 48 h with *Tug1* KD compared to both the control LNA (Figure 1B) and a non-transfected control (no LNA, Supp Fig S1A). This effect was also independently confirmed in mouse and rat skeletal muscle cells lines (Supp Fig S1B).

**Figure 1.**
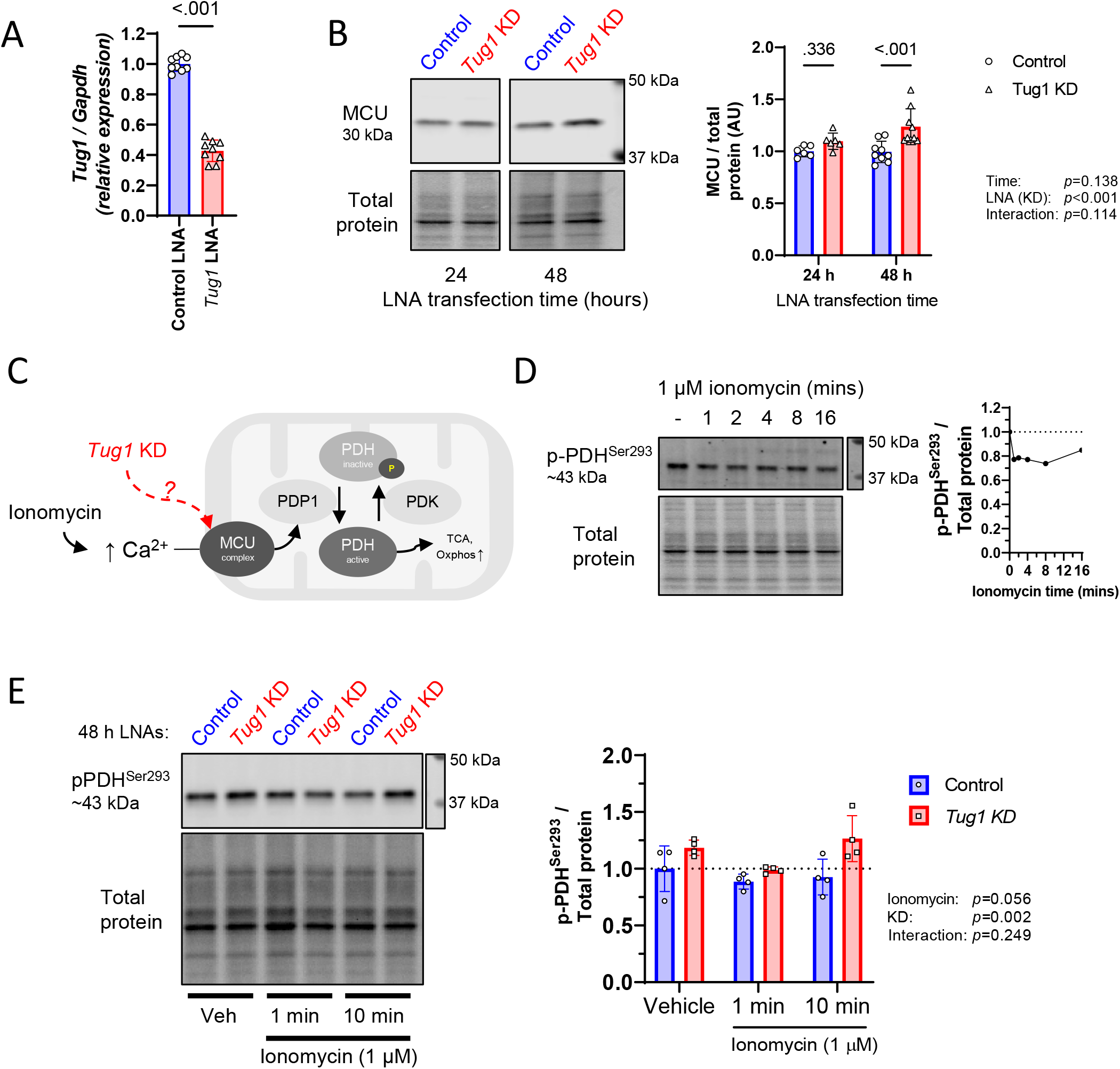
*Tug1* knockdown increases MCU expression but not a marker of Ca^2+^ uptake in H9c2 cardiomyocytes. **A)** Antisense locked nucleic acid (LNA) oligo mediated *Tug1* knockdown (KD) of RNA levels compared to a control LNA measured by qPCR. **B)** MCU protein abundance following 24 or 48 h of *Tug1* KD compared with control. **C)** Schematic depicting regulation of pyruvate dehydrogenase (PDH) via Ca^2+^-sensitive pyruvate dehydrogenase phosphatase (PDP1)-mediated dephosphorylation of PDH at Ser293. **D)** Time course of PDH Ser293 dephosphorylation in response to ionomycin (1 µM), *n*=1. **E)** PDH Ser293 phosphorylation in response to vehicle control, and 1 or 10 minutes ionomycin stimulation in *Tug1* KD compared with control cardiomyocytes. Data are mean(SD) for *n*=4 independent experiments, two-way ANOVA with Tukey post hoc test for significant main effects.

Mitochondria calcium uptake plays an important role in buffering cytosolic Ca^2+^ and is also a critical regulator of mitochondrial respiratory activity via stimulation of Ca^2+^ sensitive TCA cycle enzymes (18). In the mitochondrial matrix, Ca^2+^ ions activate PDH phosphatase 1 (PDP1), which dephosphorylates the serine 293 residue of PDH, relieving its inhibition, thus increasing PDH activity and substrate flux entering the TCA cycle (Figure 1C) (19, 20). We first characterised the phosphorylation status of PDH as a marker of mitochondrial Ca2+ uptake in cardiomyocytes over a brief time course (0, 1, 2, 4, 8 and 16 minutes) of ionomycin treatment, which increases intracellular Ca^2+^ levels (21). As expected, there was a rapid (≤1 minute) and sustained dephosphorylation of PDH^Ser293^ (Figure 1D).

Given that MCU overexpression is associated with greater mitochondrial Ca^2+^ uptake and lower PDH phosphorylation (9), we hypothesised that increased MCU with *Tug1* KD would have a similar effect and allow for a greater magnitude of PDH dephosphorylation in response to Ca^2+^ uptake. One minute of ionomycin stimulation led to a ∼12% reduction in p-PDH^Ser293^ in control cells, and ∼20% decrease in *Tug1* KD cells, although this did not reach statistical significance (ionomycin main effect p=0.056, Figure 1E). There was an overall effect for phospho-PDH^Ser293^ to be greater with *Tug1* KD (LNA main effect p=0.002; Figure 1E). After 10 minutes ionomycin, p-PDH^Ser293^ remained low in control cardiomyocytes, whereas in *Tug1* KD cells this returned to baseline levels, suggesting less mitochondrial Ca^2+^ uptake had occurred. Taken together, these data suggest that although *Tug1* KD increases MCU expression, this may not lead to a functional increase in mitochondrial Ca^2+^ uptake. The more inhibited state of PDH with *Tug1* KD may indicate that the MCU complex is less active or sensitive to Ca^2+^. This may also reflect the cell’s attempt to compensate for a disruption to the overall calcium handling network.

### Tug1 KD affects calcium pathways in the cardiomyocyte transcriptome

To gain an overall perspective of the effect of *Tug1* KD on the cardiomyocyte transcriptome, we subjected control and *Tug1* KD H9c2 cardiomyocytes to RNA-sequencing. This revealed differential expression (DE) of 1243 genes with *Tug1* KD compared with control (Figure 2A; log2fc ≥1/≤-1; FDR<0.01 and Supplemental Table S1). Gene expression of *Pdha1* and *Pdp1* were not different, but expression of *Pdk* isoforms 2 and 3 were increased with *Tug1* KD (Figure 2B), which may have contributed to the greater basal p-PDH levels (Figure 1E). Of the MCU-associated genes, the expression of *Mcu* was upregulated as expected, as was the regulatory subunit *Micu3*, and there was a small decrease in *Micu1* at FDR 0.04 (Figure 2B). The relative stoichiometry of these subunits can influence MCU sensitivity to Ca^2+^, which may have contributed to the altered p-PDH response to ionomycin with *Tug1* KD. Thus, we wanted to understand whether *Tug1* KD also affects Ca^2+^ signalling more broadly in the cardiomyocyte. Gene set enrichment analysis revealed that a number of pathways were significantly impacted by *Tug1* KD, including upregulation of “calcium regulation in the cardiac cell” (Figure 2C). Numerous genes encoding Ca^2+^ handling/signalling proteins within this gene set including *Cacna1c* (aka LTCC/DHPR/Ca_V_1.2), *Camk2g* and *Camk2b* were upregulated (Figure 2D) as were other related Ca^2+^ genes *Ncx, Pmca* and *Ryr1* (Supplemental Table S1), while *Camk2a* (Ca^2+^/calmodulin-dependent protein kinase II alpha) were downregulated with Tug1 KD (Figure 2D) along with others including the calmodulins (*Calm1-3*, Supplemental Table S1). Thus, *Tug1* KD may increase the rate of calcium cycling in the cardiac myocyte, as indicated by KEGG pathway analysis which suggested cellular uptake and intracellular of levels Ca^2+^ could be increased due to increased expression of genes encoding CaV1/2/3, along with an increased efflux due to greater expression of PMCA & NCX (Figure 2E). SR Ca^2+^ release could also be increased with *Tug1* KD due to increased expression of RyR, along with increased SR uptake due to reduced expression of PLN which would release inhibition on SERCA to increase its activity (Figure 2E). Moreover, transcription factor enrichment analysis derived from all DE genes predicted that CREB1 (CAMP responsive element binding protein 1), a downstream target of Ca^2+^ signalling via CaMKII, was the second most highly enriched among upregulated genes with *Tug1* KD (Figure 2F). Collectively, this demonstrates that *Tug1* exerts transcriptional regulation of calcium handling genes, and this may have consequences for how cardiomyocytes respond to calcium signals.

**Figure 2.**
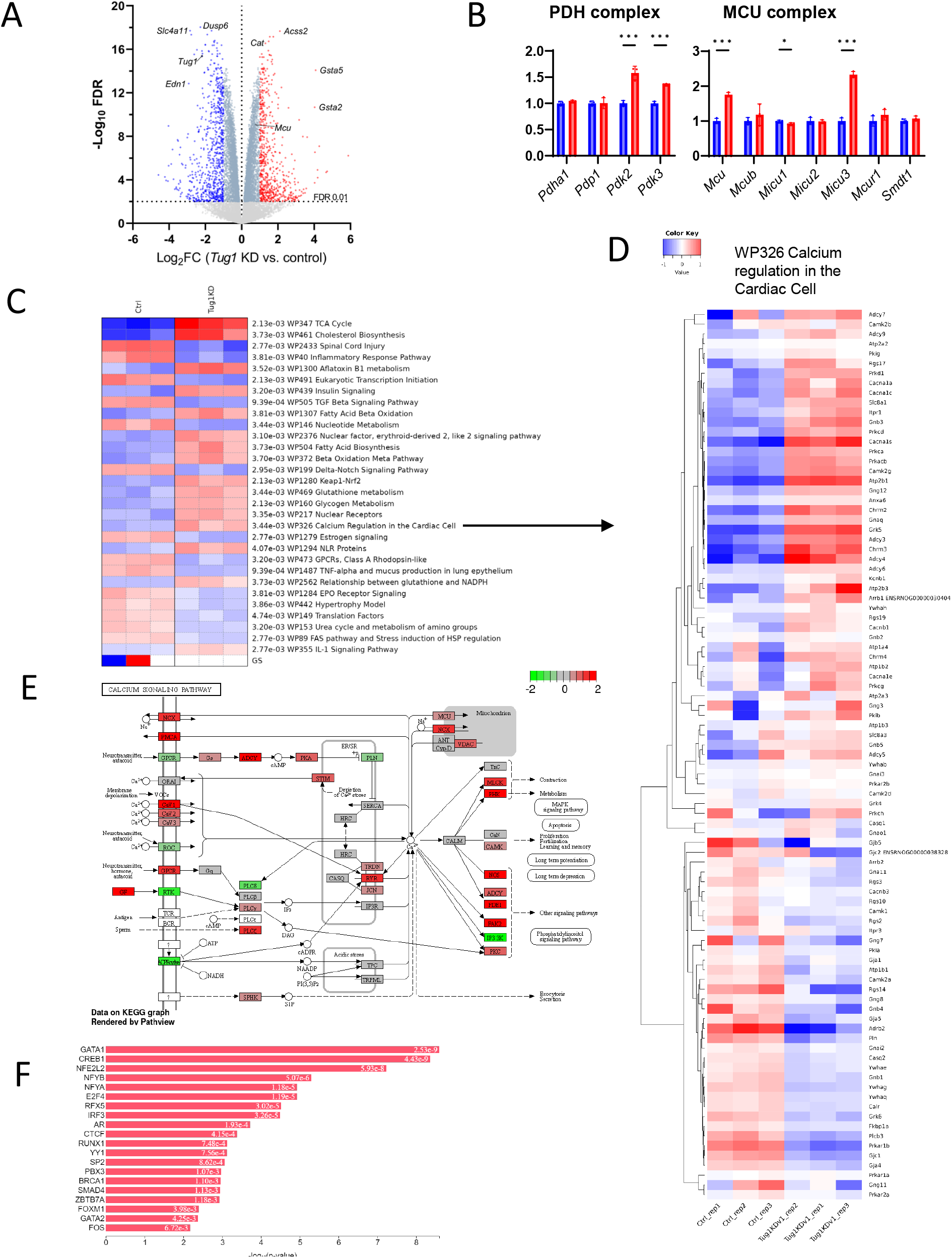
*Tug1* KD affects calcium pathways in the cardiomyocyte transcriptome. **A)** Volcano plot depicting RNA-sequencing data for *Tug1* KD compared with control H9c2 cardiomyocytes (*n*=3 replicates per group). Significantly downregulated (blue points) or upregulated (red points) at the FDR 0.01 level with log2 fold-change ≥1. **B)** Expression of genes that encode the PDH and MCU complexes, values are normalised to counts per million. **C)** Heatmap of significantly enriched gene pathways (WikiPathways, PGSEA) with *Tug1* KD compared with control. **D)** Heatmap of genes included in the “*WP326 Calcium Regulation in the Cardiac Cell*” pathway (WikiPathways). **E)** Schematic depicting up- (red) or down- (green) regulated components of the calcium signalling pathway (KEGG) based on DE genes from RNA-seq. Colour scale is log2 fold-change. **F)** Transcription factors significantly enriched (ChEA) among upregulated genes (FDR <0.05, log2fc >0.5) with *Tug1* KD compared with control.

### Tug1 KD impairs signalling responses to cytosolic C^a2^+

We next sought to investigate whether the altered expression of calcium handling genes due to *Tug1* KD leads to dysregulation of calcium signalling. CaMKII is autophosphorylated in the presence of elevated intracellular Ca^2+^, leading to activation of downstream signalling cascades involved in Ca^2+^ handling and activation of transcription factors (22). A key downstream transcription factor is the cAMP response element-binding protein (CREB), which is phosphorylated at Ser133 by CaMKII (23). To understand whether *Tug1* plays a role in this, we first characterised a time course for the response to ionomycin-induced Ca^2+^ influx in control cardiomyocytes. There was robust CaMKII phosphorylation at Thr286 within 1 minute and peaked 2 to 4 minutes (Figure 3A). We also observed the expected sequential response with the onset of maximal CREB Ser133 phosphorylation occurring at 8-16 min (Figure 3A). We then asked whether this signalling response is altered by *Tug1* KD. We found that *Tug1* KD attenuated the increase in Ca^2+^ dependent phosphorylation of CaMKII at Thr286 in response to Ca^2+^ influx by ionomycin (Figure 3B). The subsequent increase in phosphorylation of the transcription factor CREB at Ser133 was also attenuated with *Tug1* KD (Figure 3B). These data suggest that *Tug1* is required for an intact signalling response to elevated calcium in the cardiomyocyte.

**Figure 3.**
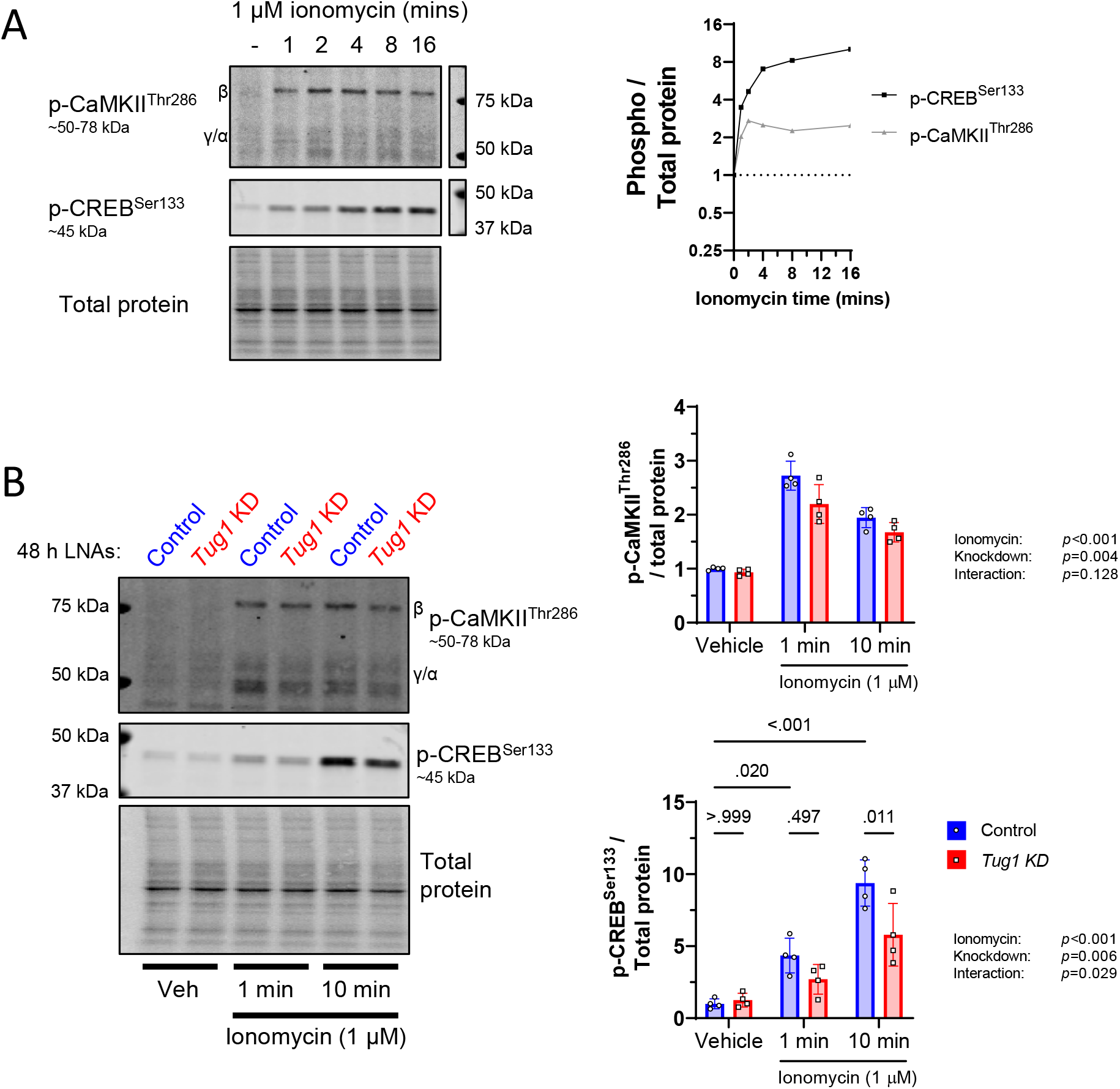
*Tug1* KD impairs cardiomyocyte signalling responses to elevated cytosolic Ca^2+^. **A)** Time course of CaMKII Thr286 and CREB Ser133 phosphorylation in response to 1 µM ionomycin (*n*=1). **B)** Phosphorylation of CaMKII Thr286 and CREB Ser133 with *Tug1* KD compared with LNA control in response to vehicle control, 1 or 10 minutes of 1 µM ionomycin stimulation. Data are mean(SD) for *n*=4 independent experiments, two-way ANOVA with Tukey post hoc test for significant main effects.

### Tug1 KD mediated MCU upregulation involves CaMKII

Finally, we sought to understand how *Tug1* KD increases *Mcu* expression. *Mcu* gene expression has been shown to be under the transcriptional regulation of CREB when phosphorylated at Ser 133 by CaMKII (24). Given that *Tug1* KD altered CAMKII and CREB phosphorylation responses to Ca^2+^ influx, and the predicted CREB involvement in the transcriptional response to *Tug*1 KD both in the present study (Figure 2E) and in our previous report in skeletal myotubes (8), we hypothesised that this pathway could be involved in the upregulation of *Mcu*. To test this, we used small molecule inhibitors of CaMKII (XII) or CREB (666-15) to investigate whether their inhibition would prevent the *Tug1* KD-mediated increase in gene expression of *Mcu*. We additionally measured a well-established CREB target gene, *Nr4a3* as a control (25). We found that the CaMKII inhibitor XII partially prevented the *Tug1* KD-mediated increase in *Mcu* expression (Figure 4A). However, the CREB inhibitor 666-15 did not prevent the *Tug1* KD-mediated increase in *Mcu* expression (Figure 4A). Both CaMKII and CREB inhibitors decreased the expression of the known CREB target gene *Nr4a3*, albeit only in the *Tug1* KD condition (Figure 4B). Collectively, this suggests that the *Tug1* KD mediated increase in *Mcu* expression occurs via a mechanism that partially involves CaMKII and a downstream step that appears to be independent of CREB.

**Figure 4.**
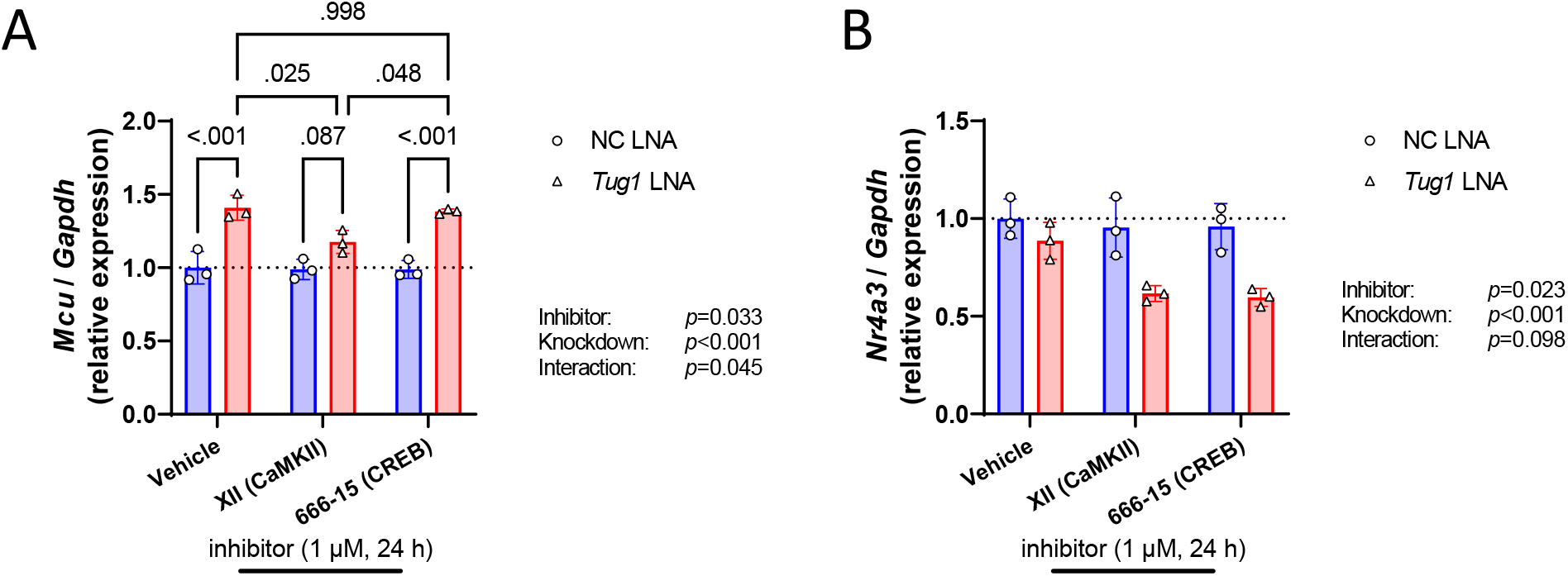
Cardiomyocyte *Tug1* KD mediated MCU upregulation involves CaMKII. Gene expression analysis by RT-qPCR of **A)** *Mcu* and **B)** *Nr4a3* in the presence or absence of CaMKII or CREB inhibitors (1 µM) throughout the 24 h control or *Tug1* LNA knockdown period. Data are mean(SD) for *n*=3 independent experiments, two-way ANOVA with Tukey post hoc test for significant main effects.

## DISCUSSION

Here we report that the long non-coding RNA *Tug1* plays a role in regulating the transcriptome in cardiomyocytes with notable effects on calcium signalling and handling. We demonstrate that *Tug1* KD leads to an increase in MCU gene and protein expression, and this event may in part require CaMKII. Despite this, the *Tug1* KD mediated increase in MCU did not appear to enhance PDH dephosphorylation, which is indicative of mitochondrial Ca^2+^ uptake.

Our finding that *Tug1* KD increases MCU expression along with numerous genes involved in calcium handling is of potential therapeutic interest, because MCU overexpression has recently been shown to reverse heart failure and rescue mitochondrial function in mice (10, 11). Despite the higher *Mcu* and *Micu3* expression with *Tug1* KD, the marker of mitochondrial Ca^2+^ uptake (PDH dephosphorylation) suggested the activity of the MCU complex was not enhanced or may have even been impaired. The MCU complex is comprised of several pore forming and structural (MCU, MCUb, MCUR1, EMRE [encoded by *Smdt1*]) and regulatory subunits (MICU1, MICU2/3) that assemble into a functional complex (5). It is possible that the extra MCU protein due to *Tug1* KD may not be incorporated into the functional MCU complex. The relative abundance of each subunit determines composition of the complex, which in turn determines the sensitivity and capacity for Ca2+ ion uptake (6). In addition, the relative expression of each subunit is tissue/cell specific, with heart having low MICU1:MICU2 ratio yet high MICU2:MCU ratio relative to muscle and liver (26). It is therefore possible that the altered relative subunit abundance with *Tug1* KD interfered with the optimal composition either within each individual MCU complex or across the population of MCU complexes.

*Tug1* KD also had a marked effect on the expression of other non-MCU related calcium pathway genes. This includes upregulation of the L-type calcium channel (LTCC; *Cacna1c*), ryanodine receptor 1 (*Ryr1*), calmodulin (*Calm1-3*) and sodium/calcium exchanger 1 (*Slc8a1*). Dysregulation of these could impact how the cardiomyocyte would respond to Ca^2+^ driven signals. In particular, the lower *Calm1-3* gene expression of calmodulin with *Tug1* KD could mean there is less calmodulin available to bind and activate CaMKII, thus potentially explaining the impaired CaMK/CREB phosphorylation that we observed in response to ionomycin. However, it remains unclear whether the differential expression of these and other calcium related genes is driven initially via *Tug1* regulating upstream transcriptional events, or whether *Tug1* affects regulation of Ca^2+^ via other downstream mechanisms, which create a feedback loop to increase expression of these calcium genes. For example, elevated cytosolic Ca^2+^ due to MCU loss of function has previously been shown to elicit broad transcriptional responses in the heart (27).

LncRNAs can elicit their regulatory effects via various mechanisms including interactions by base-pairing with RNA or DNA or with specific proteins based on lncRNA secondary structure (7). Indeed, it was reported that there are ∼3000 putative *Tug1* binding sites in the mouse genome (28). Given this, the large number of DE genes with *Tug1* KD in the present study (>1000) is not unexpected. Although it is well established that *Tug1* functions as a lncRNA (28-30), it also encodes a small open reading frame that has been shown to translated into a microprotein both *in vitro* (30) and in the human heart (31). To what extent this microprotein contributes to *Tug1*’s regulatory effects at endogenous levels remains unknown. Taken together, it is likely that *Tug1* has pleiotropic effects that should be explored in future studies.

MCU expression is regulated by a Ca^2+^/CaMKII/CREB signalling axis (9, 24). Here, we provide evidence that CaMKII is involved in the *Tug1* KD-dependent increase in *Mcu* expression. This finding was made with an inhibitor (XII) reported to be 100-fold more specific for CaMKII than the commonly used KN93 inhibitor (32). Inhibition of the downstream transcription factor CREB using 666-15 did not prevent the *Tug1* KD induced *Mcu* upregulation, although it partially blocked another CREB target gene *Nr4a3* (albeit only when *Tug1* was knocked down). A possible explanation for the lack of CREB involvement in *Tug1* mediated regulation of *Mcu*, is that CaMKII can activate a number of downstream transcription factors in addition to CREB (25). Thus, if *Tug1* regulates the activity of one or more of these TFs, this could explain the divergent expression patterns of *Mcu* and *Nr4a3* in the presence of the CaMKII and CREB inhibitors.

In summary, we show that loss of *Tug1* leads to dysregulation of the calcium handling network in the cardiomyocyte. This may have implications for understanding heart disease and the design of new therapeutic strategies, since calcium handling abnormalities are involved in the progression of heart failure, arrythmias and cardiomyopathies (1). Given that targeting calcium handling specifically at the mitochondrial level has shown promise as a therapeutic strategy in preclinical models of heart failure (9-11), further investigation is warranted into the role of *Tug1* in the heart.

## SUPPLEMENTAL MATERIAL

Supplemental Figs. S1A–S1B: DOI. https://doi.org/10.6084/m9.figshare.23730675

Supplemental Table S1: DOI. https://doi.org/10.6084/m9.figshare.23730654

## ACKNOWLEDGMENTS

We acknowledge Mark Richardson (Charles River Laboratories) for assistance with RNA-seq data pre-processing.

## GRANTS

This study was supported by a Faculty of Health Research Capacity Building Grant, Deakin University to AJT. KLW is supported by a Heart Foundation Future Leader Fellowship (102539).

## DISCLOSURES

The authors have no conflict of interest to declare.

## DATA AVAILABILITY

RNA-sequencing data from this study have been deposited at the NCBI Gene Expression Omnibus (accession number GSE237787).

## AUTHOR CONTRIBUTIONS

Conceived and designed research: AJT, GDW, SL.

Performed experiments: AJT.

Analyzed data: AJT.

Interpreted results of experiments: AJT, KLW, GDW, SL.

Prepared figures: AJT.

Drafted manuscript: AJT.

Edited and revised manuscript: AJT, KLW, GDW, SL.

Approved final version of manuscript: AJT, KLW, GDW, SL.

## Notes

https://doi.org/10.6084/m9.figshare.23730675

https://doi.org/10.6084/m9.figshare.23730654

### Competing Interest Statement

The authors have declared no competing interest.

